# Monitoring transgenic mosquitoes using wing measurements and I3S Classic

**DOI:** 10.1101/645028

**Authors:** Nayna Vyas-Patel

## Abstract

Stress responses in insects can manifest as changes in size, shape and symmetry of the wings. Developing methods to measure and track such features could act as an early warning indicator of adverse events or, if all is well, provide assurance that field or laboratory colonies were fit, healthy and developing optimally. This is especially important in the case of newly developed transgenic insects, to assess morphology and as an indicator of their fitness. As body size and symmetry is known to be a significant correlate of fitness, the potential of transgenic insects is reflected in their phenotypic expression. Microsoft Paint and Photos as well as I3S Classic were used. The wings of transgenic mosquitoes DSM 1 & 2 were measured and compared to those of the parent population *Anopheles gambiae* G3. The right and left wings of both sexes were assessed to determine if they were symmetrical. Measurements indicated high wing symmetry in all the groups and sexes tested, indicating that the transgenic mosquitoes should be just as functional as their parents. The transgenic mosquitoes DSM 1 & 2 were found to be significantly larger in length and width than the parent population *A. gambiae* G3 and could be distinguished from the parent strain using I3S Classic software with 70 to 100% accuracy. I3S Classic ranked the correct sex of the test strain predominantly in the initial ranks indicating the differences in architecture of male and female wings. I3S Classic software was also used to assess wing symmetry. In keeping with the data from taking measurements, the software indicated that the wings were highly symmetrical, both the right and left wings of the correct strain were selected in the early first and second ranks in roughly equal measure. The importance of assessing the morphological characteristics of insects and of taking measurements during the investigative procedure was discussed.

## Introduction

The creation of newly engineered organisms deserves closer phenotypic scrutiny, especially in the case of smaller sized dipteran insects such as mosquitoes. Easy to use morphological methods need to be developed to monitor their development and if differences exist, to distinguish them from closely resembling parent populations and other field strains. Physiological size, shape and symmetry are closely linked to function in insects. For example, it is known that mosquito size plays an important role in their mating preferences; comparing the size of a transgenic mosquito with that of the parent or field strain could provide valuable information about their predicted function. Some species of mosquitoes prefer to mate with females that are larger than average (Helinski MEH & Harrington LC, 2011), in others, larger males were reported as being more successful in mating than smaller ones (Yuval and Bouskila 1993; Yuval, Wekesa and Washino 1993), whilst a different study reported species that prefer to mate with individuals of a similar size to themselves (Callahan AC et al, 2018). Hence, any genetically modified mosquito needs to at least match those preferences.

The mosquitoes used here were engineered from the *Anopheles gambiae* G3 strain reared at Imperial College, London (Hammond A et al, 2016). Whilst studies have found no effect of male body size in *Anopheles gambiae* s.s. mating success (Charlwood A et al, 2002), it was found that larger females of *A. gambiae* s.s. were preferentially selected for mating by males (Okanda FM et al, 2002). In *A. gambiae*, female fecundity rapidly increased proportionally with body size (Maiga H et al, 2012). Lyimo EO & Takken W (1993) concluded that wing lengths of 3mm were the critical size permitting oviposition from the first blood meal and that there was a positive correlation between fecundity and wing-length. These findings further indicate the importance of keeping track of size in transgenic *A. gambiae* produced in the laboratory and intended for field release.

More than any other character, wings are the most reliable predictors of mosquito size and weight (Petersen V et al 2018). Larger wings indicate larger size and nutritional store, with an attendant competitive advantage. It was prudent therefore to monitor the size of transgenic mosquitoes using the wings, especially in comparison to the parent and at the same time ensure that both the right and left wings were of similar size and shape, in other words, bilaterally symmetrical.

Wings that are bilaterally symmetrical are positive indicators of functionality and fitness, asymmetrical wings would be disadvantageous in flight for obvious reasons, not least because as well as mating on surfaces, many mosquito species mate on the wing whilst flying, where compromises in aerial agility would be detrimental. Evolutionary biologists have stated that symmetry is an indication of ‘good genes’ (Scheib JE et al, 1999), others have argued that symmetry could merely be due to the lack of exposure to stressors during development (Grammer K & Thornhill R, 1994). The insertion of fluorescent markers during the formation of transgenic mosquitoes in this study (Hammond A et al, 2016) could be considered to be a stressor, it was logical therefore to examine if this or any other process during the creation of transgenic strains affected wing symmetry. Unlike other genetic experiments that directly targeted the wings to result in asymmetry (Ray et al 2016), the genes targeted to produce the transgenic mosquitoes in this study were not expected to affect the wings; the only concern was the effect of any stressors on the newly created mosquito strain.

Size, shape and symmetry can be ascertained by measuring the length and width of pairs of right and left wings and by comparing the measurements using simple mathematical tools. Image recognition software such as I3S Classic (Hartog JD & Reijns R, 2013) could also be used to assess how closely the transgenic mosquitoes resembled the parent strain. As wing shape was annotated in I3S when wing outlines were marked where the veins met the rim of the wing, using the software could take any shape changes of the wings into account (Vyas-Patel N & Mumford JD 2017). To that end, the wing length and width of thirty five male and female transgenic Dominant Sterile Male Strain 1 & Strain 2 (DSM 1 & 2) as named by Target Malaria (Valerio L et al, 2016; Target Malaria, 2019) and the parent strain *A. gambiae* G3, were measured using Microsoft Paint and the right and left wings compared using measurements and the software, I3S Classic.

## Method

The right and left wings of transgenic DSM 1 & 2 and *A. gambiae* G3, of both sexes were dissected out under the microscope and each pair of right and left wings were photographed using a Sony Cybershot DSC-W800 camera. The photographs were all taken at a set camera magnification of three, on a digital setting, before measurement in Microsoft Paint; this is also known as Paintbrush and Windows Paint and is free to download.

Thirty five male and female transgenic DSM1 & 2 and *Anopheles gambiae* G3, were examined. Both the right and left wings were photographed and the images were used for measurement and later in I3S Classic.

Twelve groups were tested, the right and left wings of both sexes of DSM 1 & 2 and of *A. gambiae* G3.

### Wing Measurements

Measurements of the length and width of the wings were taken as shown in Figure 1. The images were opened using Microsoft Paint and the units in Paint were calibrated to the camera magnification. It was found that five units on the Paint ruler were equivalent to 6.8 cm, hence each Paint unit was equivalent to 1.36 cm. However, Paint had been set to the cm scale, so the results were divided by 10 to obtain the correct measurements in mm. This calibration was used throughout. All of the measurements for both sexes and for the length and widths for each group were examined using analysis of variance (ANOVA), Tables 1 and 2. Box and line plots of the ANOVA results were further examined, Figures 2, 3, 4 and 5. The measurement for the right and left wings were compared using the Student’s t test, Table 3. The mean, mode and the median for the right and left wings were individually examined graphically for each group, Figure 6. The mean, the mode and the median values (measures of central tendency) were placed together for each group for the right and left wings and were presented and examined graphically, Figure 7. The measurement for the right wings were subtracted from the corresponding left wings (Table 4) to determine if there was a large difference from each pair of wings.

### Comparison of the Right and Left wings using I3S

Thirty five wing images of both sexes of DSM1 and 2 and *Anopheles gambiae* G3 were marked in I3S Classic (free to download) as described previously by Vyas-Patel N & Mumford JD (2017 & 2018). Wing images were aligned as in Figure 1 using Microsoft Photos (free to use and download). Aligned images were uploaded into I3S and marked as before (Vyas-Patel N & Mumford JD, 2017 & 2018). One image each of the right and left wing of both sexes in each group were uploaded into the I3S database for testing. As the database contained an image of the right and left wings of both sexes from each group, a total of 12 wing images were present in the I3S database. Each of the 35 marked wings in every group was tested in I3S and the results were noted up to rank 12 (there were 12 wings in the database). A sample of the rankings were presented in Tables 4 and 5. For the rest of the groups, the trends in the results were outlined and the total percentages of correctly retrieved results at rank 1 were stated beneath Tables 4 and 5.

## Results

**Figure 1:**
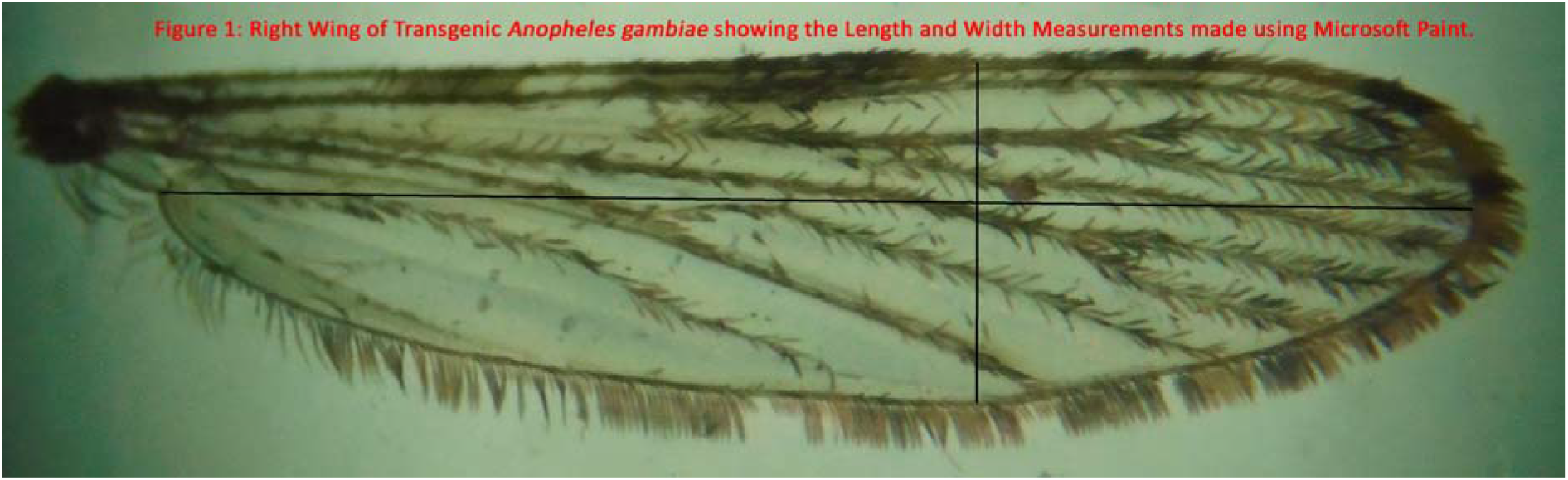
Indicating the points used on the wing for the length and width measurements in Microsoft Paint.

The length was measured using the kink at the point of insertion of the wing into the thorax on the left (alular notch), as the starting point and ending at the furthest point on the curve of the wing on the right. The width was measured using the widest points on the wings, starting just before the wing curved up from the bottom edge of the wing.

**Table 1:** Analysis of Variance (ANOVA) results performed on the Length Measurements.

| Source of Variation | Sum of Squares | Degrees of Freedom, d.f. | Mean Squares | F Value |
| --- | --- | --- | --- | --- |
| Between Groups | 3.878 | 11 | 0.3535 | 25.94 |
| Error | 5.545 | 408 | 1.3592E-02 |  |
| Total | 9.423 | 419 |  |  |

The probability of the above result, assuming the null hypothesis, was less than 0.0001. The p value was calculated using an online calculator - https://www.socscistatistics.com/pvalues/fdistribution.aspx.

For an ‘F’ value of 25.94 and d.f. of 11 (between treatments) and d.f. of 34 (within treatments), the p-value was <0.00001. The result was significant at p <0.05.

**Table 2:** Analysis of Variance (ANOVA) results performed on the Width Measurements.

| Source of Variation | Sum of Squares | Degrees of Freedom, d.f. | Mean Squares | F Value |
| --- | --- | --- | --- | --- |
| Between Groups | 1.215 | 11 | 0.1104 | 110.0 |
| Error | 0.4096 | 408 | 1.0039-03 |  |
| Total | 1.624 | 419 |  |  |

The probability of the above result, assuming the null hypothesis, was less than 0.0001. The p value was calculated using an online calculator - https://www.socscistatistics.com/pvalues/fdistribution.aspx.

For an ‘F’ value of 110.0 and d.f. of 11 (between treatments) and d.f. of 34 (within treatments), the p-value was <0.00001. The result was significant at p <0.05. Differences were present in wing length and width measurements between groups.

**Figure 2.**
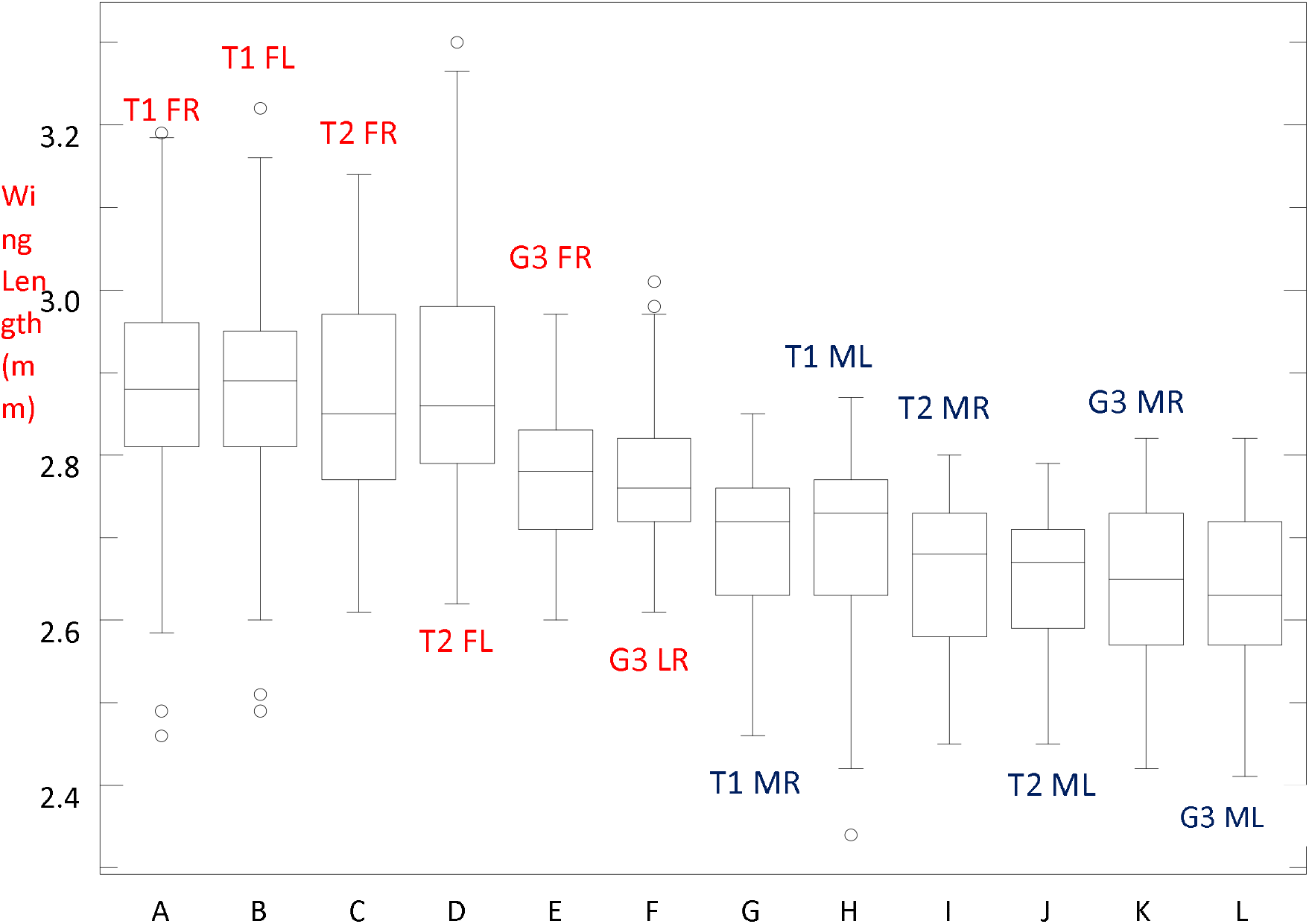
Box Plot (ANOVA) of Wing Lengths. T1 & 2 = DSM1 & 2; G3 = Parent strain (*A. gambiae* G3); R= Right wing; L = Left Wing; F= Female; M = Male.

In all cases (female and male mosquitoes, right and left wings), the parent *A. gambiae* G3 strain had lower length measurement values than both the transgenic strains DSM 1 & 2, indicating that both DSM1 & 2 transgenic strains were longer in length than the parent strain, G3. As there were a large number of replicates (35 in each group), the outliers did not contribute significantly to the results – the trends were still clear. Any skew within the box plots, away from the median values should not be viewed to be significant as it is more than likely to arise from the operator carrying out all of the tasks involved in wing measurements. Values for male wings were much lower than female wings in all groups, reflecting the known smaller size of males compared to female *A. gambiae* mosquitoes. This trend continued in the transgenic mosquitoes – male DSM 1 & 2 had lower values (and hence were smaller) than female DSM 1 &2. Transgenic females were longer in length than transgenic males (DSM 1 & 2) as well as the parent G3 strain. Female G3s were longer than male G3s and also longer than the transgenic males.

Unpaired t tests comparing T1 & T2 Female (right wings P = 0.8521; left wings P = 0.8233) and T1 & T2 Male (right wings P = 0.124; left wings P = 0 1864); indicated that there was no significant difference between T1 & T2 (DSM 1&2) transgenic wing lengths (P>0.05).

**Figure 3.**
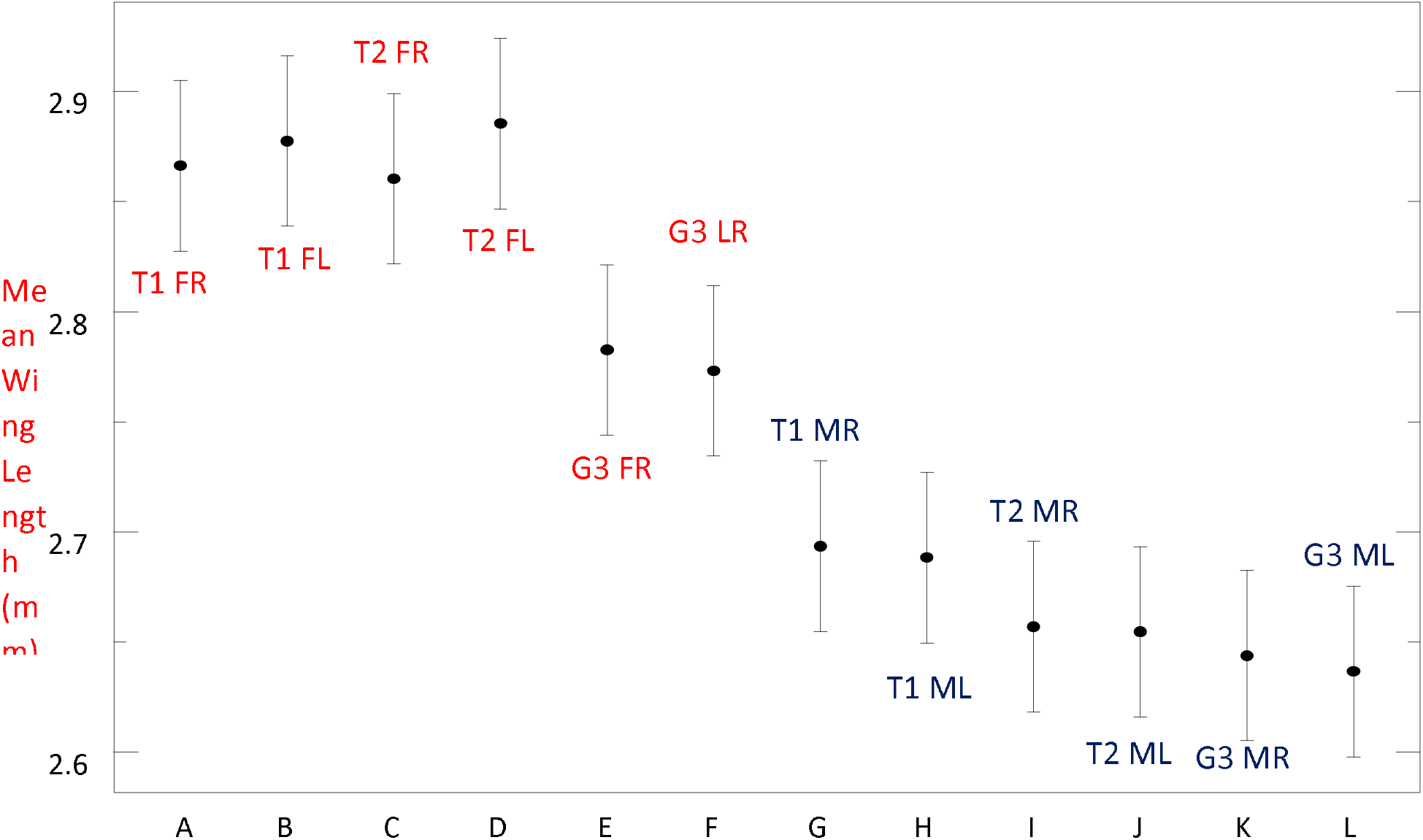
ANOVA Mean Wing Lengths. T1 & 2 = DSM1 & 2; G3 = Parent strain (*A. gambiae* G3); R= Right wing; L = Left Wing; F= Female; M = Male.

The *A. gambiae* G3 males were the shortest and the transgenic females were the longest. Both sexes of transgenic mosquitoes were longer than the parent G3 strain.

**Figure 4.**
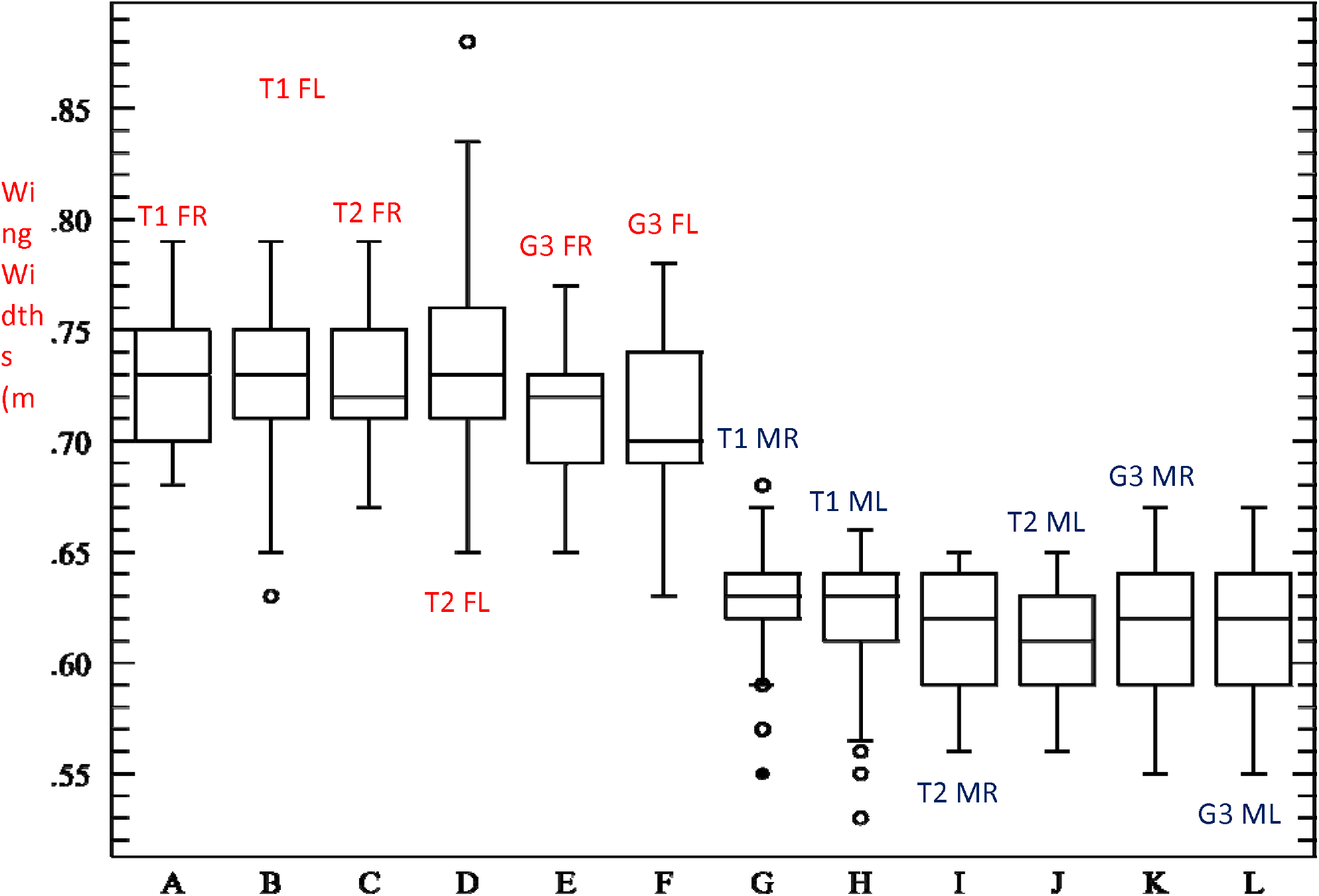
Box Plot (ANOVA) of Wing Widths. T1 & 2 = DSM1 & 2; G3 = Parent strain (*A. gambiae* G3); R= Right wing; L = Left Wing; F= Female; M = Male.

The range – the difference between the maximum and minimum indicated by the ends of the lines from the boxes (the whiskers) and the outliers (the circles), varied amongst the groups. This may largely be due to the fact that the width was not constant across the wing and even though measurements were taken at the widest point of the wing, this was by eye and could contribute to the ranges seen. Furthermore, despite the weight of the coverslip, wings could still be slightly folded between the veins, contributing to the larger range in results. The median values (horizontal lines within the boxes) were largely constant.

Unpaired t tests comparing T1 & T2 Female (right wings P = 0.9074; left wings P = 0.5673) and T1 & T2 Male (right wings P = 0.0654; left wings P = 0 0713); indicated that there was no significant difference between T1 & T2 (DSM 1&2) transgenic wing widths (P>0.05).

**Figure 5.**
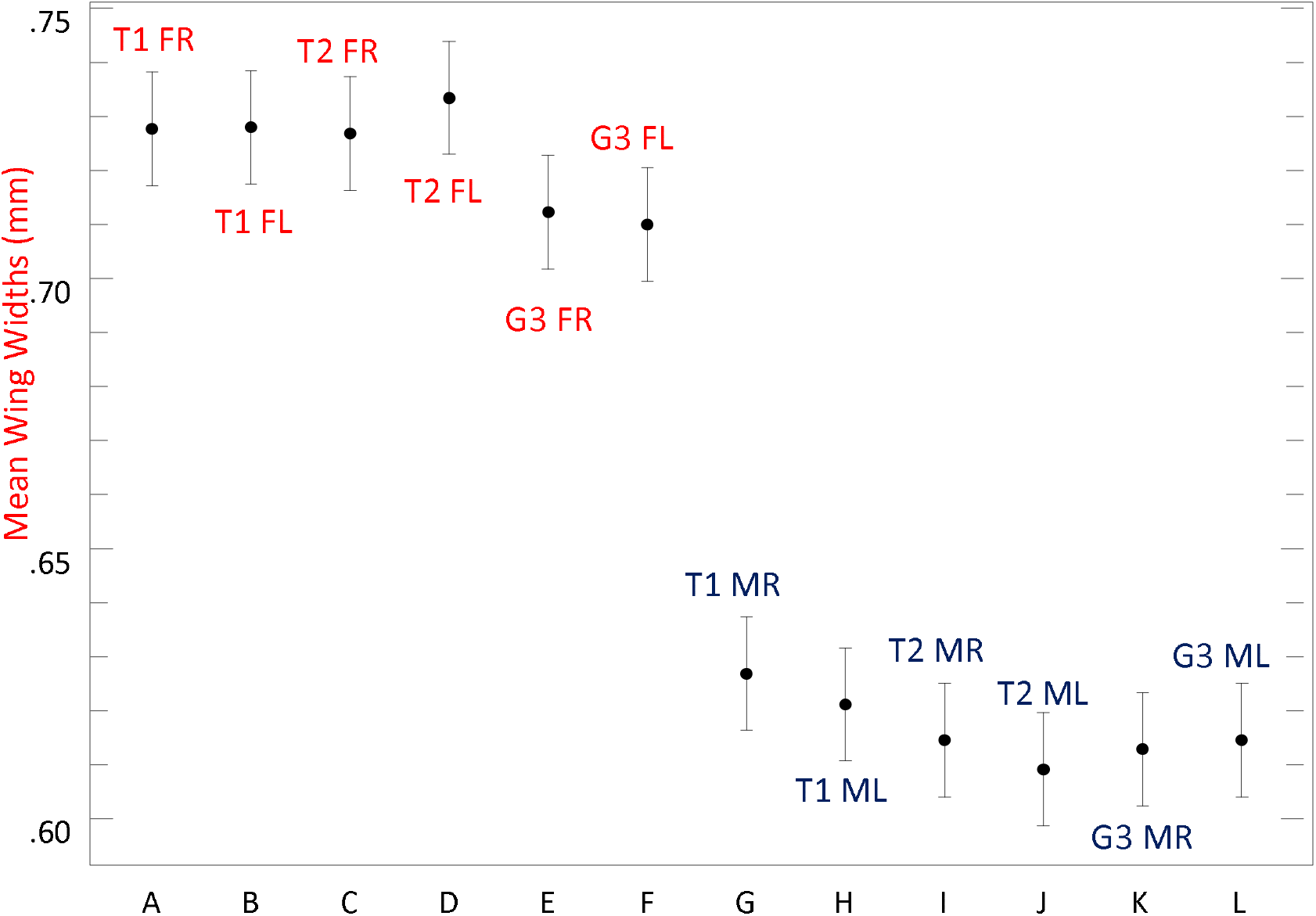
ANOVA Mean Wing Widths. T1 & 2 = DSM1 & 2; G3 = Parent strain (*A. gambiae* G3); R= Right wing; L = Left Wing; F= Female; M = Male.

Comparisons of widths reveal trends similar to length measurements. Transgenic mosquitoes (DSM 1 & 2) were larger and broader than the parent stock (*A. gambiae* G3), especially in the case of females. Male wings were visibly demarcated from the females by being narrower in width, across all the groups, in both transgenic and parent mosquitoes. This indicated that wing widths could also be used as a reliable indicator of the sex of a mosquito, with narrower wing widths indicating male mosquitoes. The Analysis of Variance was calculated using The College of Saint Benedict and Saint John’s (CBSJU) online ANOVA calculator (http://www.physics.csbsju.edu/stats/anova.html). The raw results may be viewed in ‘Supplementary Material’.

**Figure 6:**
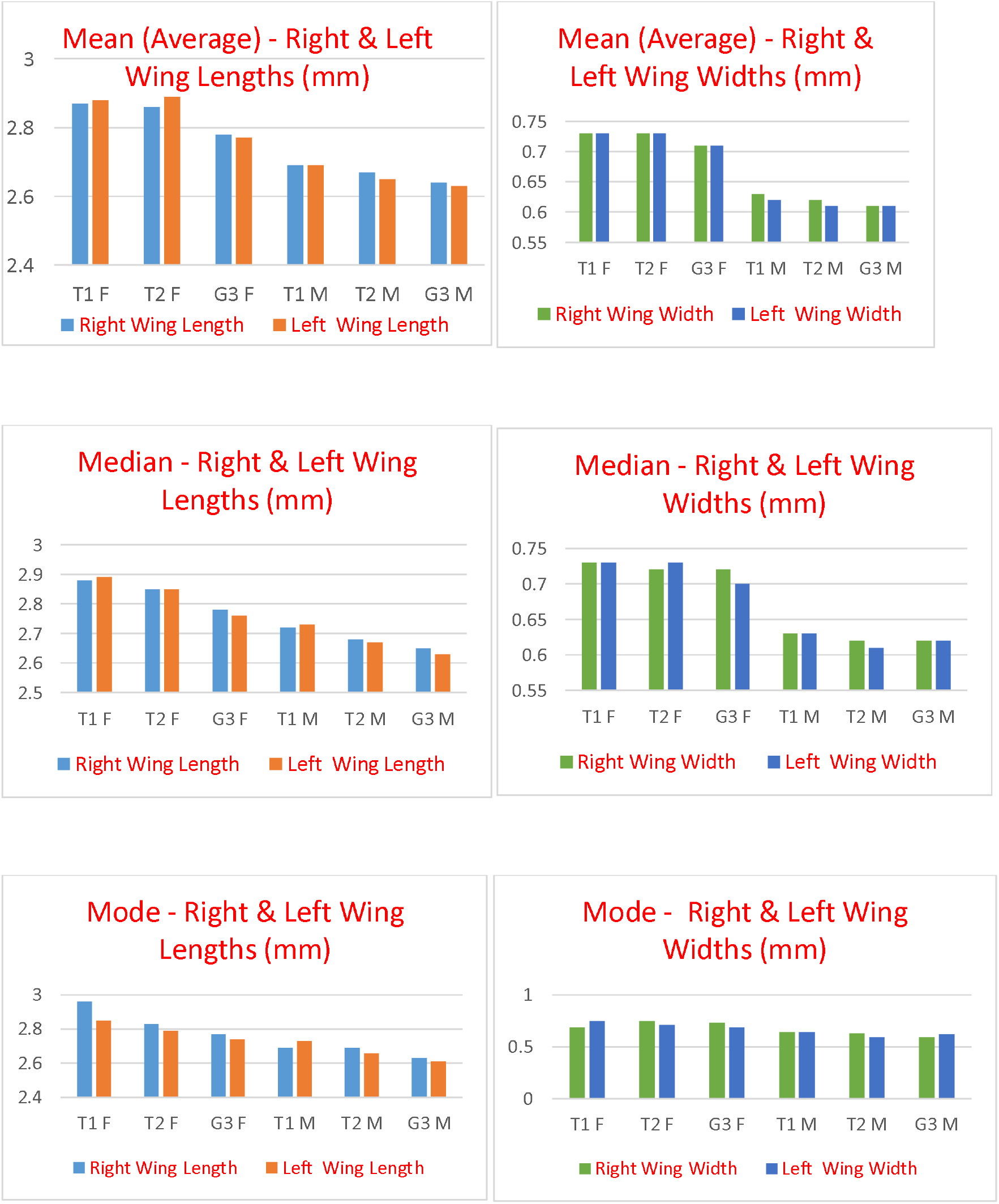
The Mean, the Mode and the Median for Right & Left Wings. F = Female wings; M = Male wings; T1 & T2 = DSM 1& 2; G3 = A. gambiae G3 strain (Parent). Very little difference exists between the middle values (Mean, Mode, Median) for right and left wings in all the groups tested.

**Figure 7:**
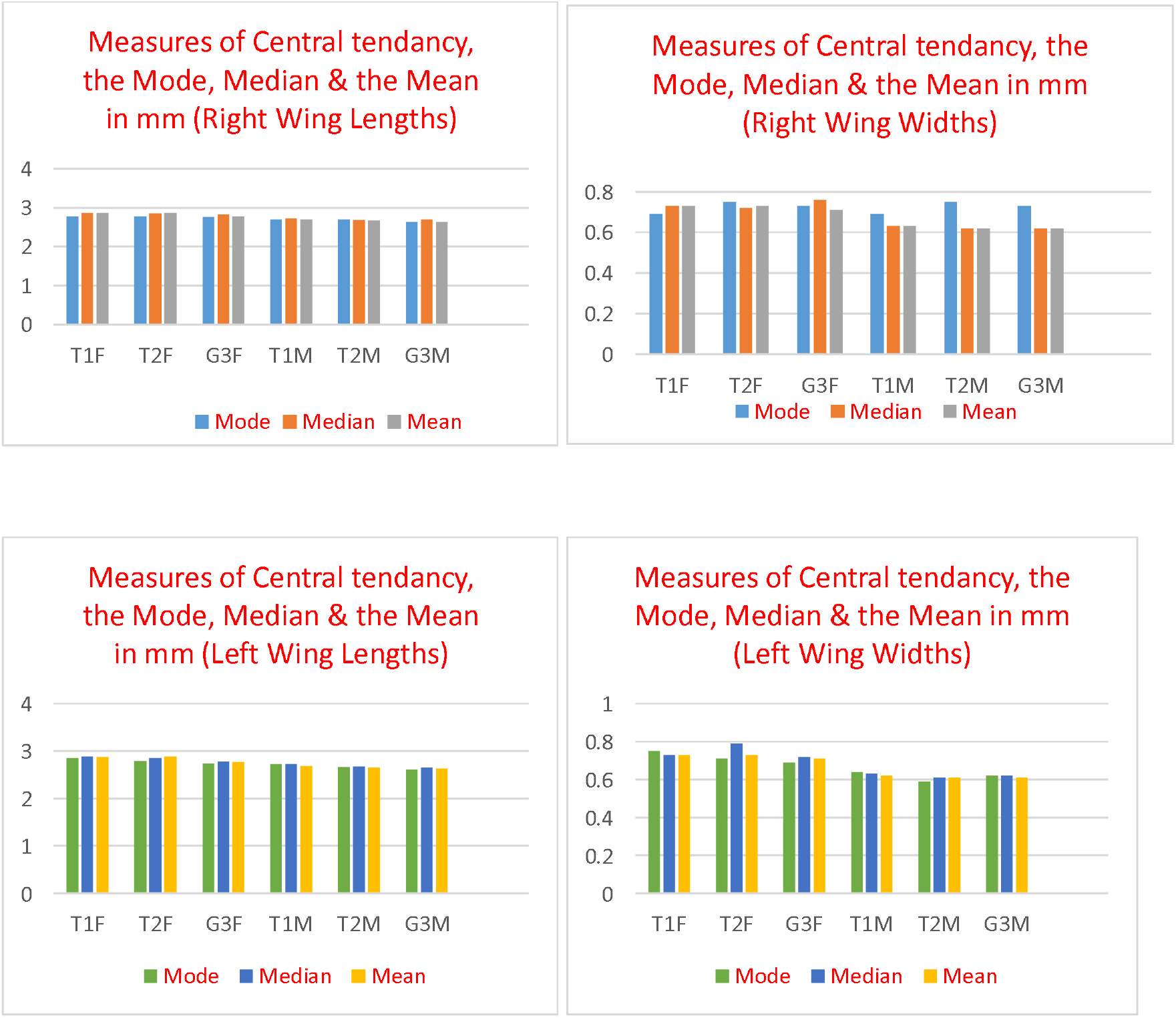
The similarity of the middle values- the Mode, Median and the Mean.

The middle values (the Mode, the Median and the Mean) for the measurement of lengths and widths can be seen to be fairly uniform and did not differ greatly between the transgenic and parent populations (Figures 5 & 6). It would be expected that in a symmetrical distribution, the mean, the mode and the median values would be close to each other as they are here. A slight variation in the widths of both right and left wings in both sexes can be seen, reflecting the fact that the width is not uniform across the lower edge of the wing and even the best efforts to locate the widest measure of the width leads to a slight variation in the middle values. This was not significant, Table 3 below.

**Table 3:** Difference between the right and left wings of DSM 1&2 (T1 &2) and A. gambiae G3 mosquitoes using a paired ‘t’ test. t = Paired t value (2 tailed); SE = Standard Error of difference; P = P value at 95% C.I. (Confidence Interval); Mean values are of the right minus left wing. The degrees of freedom (d.f.) in all cases was 34. The ‘t’ test results were calculated using Graph Pad’s Quickcalcs online calculator (https://www.graphpad.com/quickcalcs/ttest1/).

|  | P value<br>Wing<br>Length | t value<br>Wing<br>Length | S.E. | Mean | P value<br>Wing<br>Width | t value<br>Wing<br>Width | S.E. | Mean |
| --- | --- | --- | --- | --- | --- | --- | --- | --- |
| T1 F | 0.2886 | 1.0780 | 0.010 | -0.0111 | 0.9385 | 0.0777 | 0.004 | -0.0003 |
| T1 M | 0.7267 | 0.3524 | 0.015 | 0.0051 | 0.2408 | 1.1940 | 0.005 | 0.0057 |
| T2 F | 0.0528 | 2.0064 | 0.013 | -0.0251 | 0.5044 | 0.6747 | 0.007 | -0.0049 |
| T2 M | 0.7467 | 0.3257 | 0.007 | 0.0023 | 0.1236 | 1.5788 | 0.003 | 0.0054 |
| G3 F | 0.3730 | 0.9028 | 0.010 | 0.0094 | 0.3996 | 0.8530 | 0.004 | 0.0031 |
| G3 M | 0.2084 | 1.2825 | 0.006 | 0.0071 | 0.2053 | 1.2913 | 0.001 | -0.0017 |

There was no significant difference between the right and left wing measurements (p> 0.05) in either the parent (*A. gambiae* G3) or transgenic mosquitoes (DSM 1, 2 denoted as T1, T2).

**Table 4:** The difference between the right and left wings in each group, average values. F = Female wings; M = Male wings; L = left Wing; R = Right Wing; T1 & T2 = DSM 1& 2; G3 = *A. gambiae* G3 strain (Parent).

| Average Values of Wing Differences | T1FR-T1FL | T2FR-T2FL | G3FR-G3FL |
| --- | --- | --- | --- |
| Female Lengths | 0.01114 | 0.02514 | 0.009429 |
| Female Widths | 0.00029 | 0.00657 | 0.002286 |
| Male Lengths | 0.005143 | 0.002286 | 0.007143 |
| Male Widths | 0.005714 | 0.005429 | 0.00171 |

The right wing values were subtracted from the left wing values (for each pair of wings) and the average for each group was presented above. The closer this value is to zero, the more symmetrical the wings are. The results indicate that the wings were highly symmetrical in each group. There were 35 individuals in each group.

**Table 5:** Ranked results indicating the demarcation of the sexes and groups when wings were tested in I3S. Results for Female 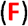 DSM1 Left and Right (T1 L & T1 R) Wings.

| Test | T1 FL | T1 FL | T1 FL | T1 FL | T1 F L | T1 R | T1 R | T1 FR | T1 FR | T1 FR |
| --- | --- | --- | --- | --- | --- | --- | --- | --- | --- | --- |
| Rank 1 | T1 FL | T1 FL | T2 FL | T1 FR | T1 FR | T1 FL | T1 FL | T1 FL | T1 FR | T1 FL |
| Rank 2 | T1 FR | T1 FR | G3 FL | T1 FL | G3 FR | T1 FR | T1 FR | T1 FR | G3 FL | T1 FR |
| Rank 3 | T2 FR | T2 FR | T1 FR | T2 FL | T2 FL | G3 FL | G3 FL | G3 FL | G3 FR | T2 FL |
| Rank 4 | G3 FR | G3 FL | G3 FR | T2 FR | T1 FL | T2 FR | G3 FR | T2 FL | T2 FR | G3 FL |
| Rank 5 | G3 FL | T2 FL | T2 FR | G3 FR | T2 FR | T2 FL | T2 FR | G3 FR | T1 FL | G3 FR |
| Rank 6 | T2 FL | G3 FR | T1 FL | G3 FL | G3 FL | G3 FR | T2 FL | T2 FR | T2 FL | T2 FR |
| Rank 7 | G3 ML | T1 MR | T1 MR | G3 MR | G3 ML | G3 MR | T1 MR | T1 MR | G3 ML | G3 ML |
| Rank 8 | T1 ML | G3 ML | T2 MR | T1 MR | G3 MR | T2 MR | T2 MR | T1 ML | G3 MR | G3 MR |
| Rank 9 | T1 MR | T2 ML | T1 ML | G3 MR | T1 MR | G3 ML | T2 ML | G3 MR | T2 MR | T1 MR |
| Rank 10 | T2 MR | T2 MR | G3 ML | T2 ML | T2 MR | T1 MR | G3 ML | G3 ML | T1 MR | T2 ML |
| Rank 11 | G3 MR | T1 ML | G3 MR | T1 ML | T1 ML | T1 ML | T1 ML | T2 ML | T1 ML | T1 ML |
| Rank 12 | T2 ML | G3 MR | T2 ML | T2 MR | T2 ML | T2 ML | G3 MR | T2 MR | T2 ML | T2 MR |

**Table 6:**
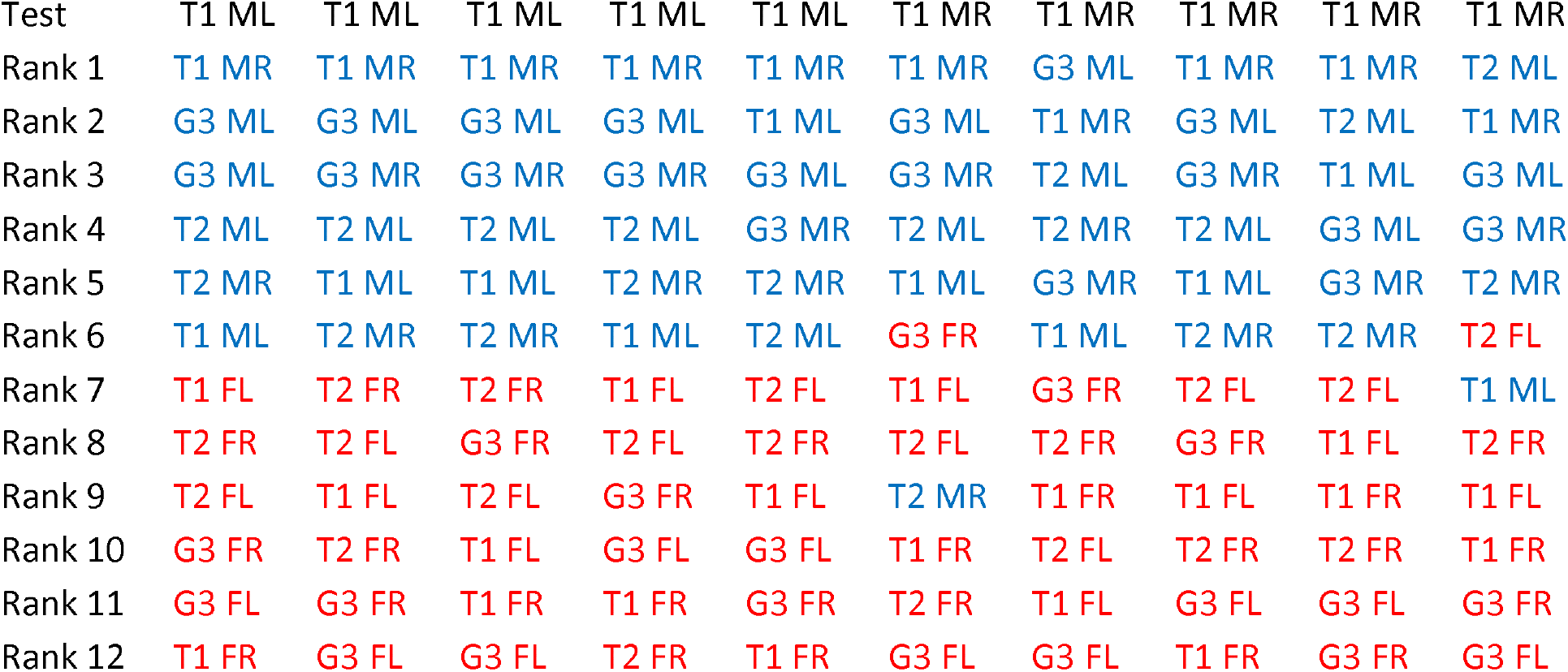
Ranked results indicating the demarcation of the sexes and groups when wings were tested in I3S. Results for Male 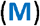 DSM1 (T1) Left and Right (T1 L & T1 R) Wings. T1 & 2 = The transgenic strains, DSM 1 & 2. G3 = *Anopheles gambiae* G3, parent strain. F = Female wing (coloured red), M = Male wing (coloured blue). R = Right wing, L= Left Wing. The test wing was either the right (T1 R) or Left (T1 L) wings of DSM 1 (T1).

Tables 4 & 5 depicts the raw results when either the right (R) or left (L) wing of female and Male DSM 1 (T1) were tested in I3S. All of the female wings were ranked 1 to 6 when tested with a female wing of DSM 1 (vice versa for male wings). The remaining wings further down the ranks were of the males from the different groups retrieved from the database. These were ranked 7 to 12. There were only 12 wing images in the database, six of each sex and if a female wing was the test, the female wings from the database were all ranked early, followed by the male wings. Vice versa when the test was of a male wing. A similar demarcation of the sexes was seen when the wings of DSM 2 and G3 were tested in I3S. In a few cases, males could be seen in later ranks six to nine when tested with a female wing and vice versa for the other sex, Table 5, but this happened infrequently (twice in the case of DSM1 & 2 and four times in the case of G3, not shown above).

Rank 1 results indicate that when tested with a right wing, the left wing was chosen just as much in the early ranks 1 &2 (if not more in the case of males) as the right one (vice versa for the left wing) demonstrating that there was little difference between the wing shapes and the pattern of the right and left wing markings. 40% of the correct sided female wings (i.e. left or right) were chosen at rank 1 in Table 5 and 30% of the male wings in Table 6. This pattern was repeated in the other cases. For DSM 2, 50% correctly sided wings were selected at rank1 for females and 40% for males. For *A. gambiae* G3s, 60% of correctly sided wings were selected at rank 1 in the case of females and 40% in the case of males. A pattern was not observed in the selection of left and right wings at rank1, the right or left wings were equally likely to be chosen at rank 1 or 2 (i.e. the early ranks).

Taking all of the results into account, at Rank 1, the percentage of correctly identified strains was 100% (T1F); 90% (T1M); 100% (T2F); 70% (T2M); 70% (G3F); 80% (G3M). I3S identified the correct strain at Rank 1, with 70% to 100% accuracy.

## Discussion

Gene modification could come with unintended consequences which need to be assessed and monitored. Before anything else is considered, phenotypic examination and measurement could reveal important information about the newly created organism. Morphology and physical dimensions (size, shape and structure) affect and are closely related to function in insects.

Mosquitoes depend on sexual reproduction for species maintenance and transference of genetic traits. When there are concerns about size assortative mating in many mosquito species (Sawadogo et al, 2013; Paton D et al, 2013; Scott TW, 2002; Lyimo EO & Takken W, 1993; Okanda FM et al, 2002), it becomes even more important to keep watch on the size of any newly created mosquito strain. Studies have shown that mosquito species have evolved to recognize different sizes in mates and that smaller size equates to poor larval growth conditions and subsequent survival and desiccation resistance, resulting in preferential mating with larger or intermediate sized mates. Size assortative mating means that a larger body size is equated with higher phenotypic quality (Nghabi KR et al 2008). Any genetic modification technique should therefore be aiming to produce individuals that were either slightly larger than or at the very least equal and comparable in size to the parent and/or the field population. Whilst wing, hence body size matters, there could be a possibility that too large a size may be deleterious to mating preferences, but the extent of what constitutes a really ‘large’ or very ‘small’ size is not known. An educated guess from this study could be if transgenic male wings were larger than the female wings of the parent or field population (here transgenic males were not larger than the parent females, Figs. 2, 3, 4 & 5), it might prove disadvantageous, but that remains a hypothesis.

The demarcation of the sexes based on size (sexual dimorphism) is generally a feature of biting mosquito species where blood feeding females are significantly larger in size than the males. In contrast, non-biting, carnivorous mosquito species such as *Toxorhyncites brevipalpis* do not present with sexual dimorphism, males and females being of similar size (Corbet PS and Ali H J, 1987). In *Toxorhycites rutilus* and *Toxorhyncites amboninensis*, the females were similar in size in the case of *T. rutilus* and only slightly larger than the males in the case of *T. amboninensis* (Lounibos LP et.al, 1996). Exceptions to this generalisation have been documented, as Lounibos LP (1995) reported an absence of wing sexual dimorphism in the malaria transmitting, hence blood feeding mosquito *Anopheles darlingi*. The author noted “This absence of sexual size dimorphism is rare among mosquitoes and has not been noted previously in the genus *Anopheles*” (Lounibos LP, 1995) In general, the reduction or absence of sexual size dimorphism in non-blood feeding and non-biting mosquitoes contrasts with blood feeding mosquito species where females are significantly larger than males in the majority of species studied. The fact that the transgenic strains always mated with the parent stock when back crossed continuously, should provide assurance that the size of the transgenic mosquitoes produced here did not deter mating, at least in the laboratory. As mosquitoes with larger sized wings have been shown to be favoured for mating in a number of blood feeding mosquito species, the larger size of the transgenic mosquitos DSM1 & 2 can only be considered as a positive trait of fitness. All of the groups were fed the same diet and reared under the same laboratory conditions. Size might alter once transgenic mosquitoes were released and had undergone a few generations in the field and this would require monitoring. In this investigation, when considering wing length and widths, the transgenic populations of DSM 1 & 2 were significantly larger than that of the parent population *A. gambiae* G3 (Tables 1 & 2; Figures 2 to 7).

Previous studies reported that their transgenic insects were generally larger than the field and laboratory population from which they were derived (Yeap LH et al, 2013; Santos MN et al, 2010). This was attributed to consistent optimum rearing and nutritional conditions available in the laboratory. Both environmental and genetic influences can cause individuals of an insect species to differ in size (Perl CD and Niven JE, 2016). It was not the aim of this study to attribute the cause of any size changes determined here, only to develop simple and easy to use methods of assessing size, shape and symmetry. In insects, size is not just a significant correlate of fitness but due to its plasticity it is adaptive, enabling growing insects to survive in environments prone to fluctuations in the quantity and quality of food available (Parker J and Johnston LA, 2006). Barfi concluded that long term colonisation of *A. gambiae* in the laboratory did not affect size (Barfi E, 2015), probably due to stable and optimum laboratory rearing conditions. Here, the transgenic mosquitoes were continually being back crossed with the parent strain over many generations, the effect of any inbreeding would need to be considered and would require further study. Developing techniques such as were explored here could help to keep track of any size changes over time.

Wing size, shape and symmetry are all of importance in assessing the potential function of insects. Of the different types of asymmetry that can occur - directional asymmetry where the development of a character occurs to a greater extent on one side of a plane than another, antisymmetry where asymmetry is the normal state for the organism but where it varies on which side it could occur and fluctuating asymmetry where symmetry is the norm for the organism but asymmetry results from an inability of the organism to develop in precisely determined pathways; it is fluctuating asymmetry that has been most examined in insect wings. The three types of asymmetry are related (Graham JH 1998, Kark S 2001) and controlled by a single gene for asymmetry and chirality named Myosin 1D (Noselli S et al, 2014, Lebreton G et al, 2018). Van Valen (1962) stated that fluctuating asymmetry at least, was “commonly used to estimate the effects of minor developmental accidents”. Wing asymmetry is usually the result of genetic or environmental stress and can be induced by targeted gene modification or infections in insects. Ray RP et al (2016) targeted RNA interference to alter wing shape beyond the norm found in natural populations of Drosophila and found that it resulted in a profound and direct effect on flight performance. They demonstrated that altering the expression of a single gene significantly altered aerial agility in Drosophila and drew some important conclusions related to performance from their findings (Ray RP et al, 2016). Such a response was not expected from the transgenic mosquitoes tested here which were engineered to produce sterility (Hammond A et al, 2016) however, the effect of any associated stress needed to be monitored. Apart from the effects of possible stress, it was felt that the size and morphology of any newly created organism should be monitored, considered good practise and be part of routine operating procedure.

In a study investigating fluctuating asymmetry as a quality control indicator in the laboratory mass production of sheep blowfly *Lucilia cuprina*, Clarke GM and Mckenzie LJ (1992) concluded that “Fluctuating asymmetry may have considerably more potential as a quality control indicator and monitoring tool than other commonly used parameters”. Maiga et al (2012) predicted male attractiveness in mosquitoes using a measure of the fluctuating asymmetry in wing length. The level of symmetry in the length of left and right wings, which were thought to be a possible cue used by females to assess male quality was quantified. No significant difference between mated and unmated males of *Anopheles gambiae* was found in Maiga’s study, the wings were bilaterally symmetrical in both groups. Similarly, no significant difference was found between the right and left wings in any of the groups tested here using different methods, they were symmetrical in all the groups-tested (Table 3 & 4; Figures 2 to 7). If size is being assessed, it is straightforward and easy to also include measurements of both the left and right wings for the added assurance of wing symmetry. The simplest measure of symmetry, where the difference between the right and left wing measurements is very close to zero (Table 4) indicated that the wings in all the groups tested were highly symmetrical.

The outcomes of wing asymmetry are varied. Insects naturally exhibiting some degree of fluctuating asymmetry were found to have compensatory mechanisms and used asymmetry in wings to facilitate manoeuvers (Haas & Carter 2008). The extent of the asymmetry was important, lower levels of asymmetry having little or no effect on flight performance compared to higher levels of asymmetry which had profound effects on the ability of insects to function optimally (Fernández MJ et al, 2017). When any fluctuating asymmetry is considered in a newly created organism, the levels of symmetry/asymmetry should be comparable and match that of the parent or that of the field population for which it was created. Comparing the right and left wing measurements using different tests (Tables 3 & 4; Figures 6 & 7) clearly indicated that there was no wing asymmetry present in either the parent or the transgenic mosquitoes tested here. When wing sizes were considered, the newly created transgenic organisms were larger in length and width than the parent strain from which they were derived (ANOVA Tables 1 & 2; Figures 2, 3, 4 & 5), this did not deter mating in the laboratory between the transgenic mosquitoes and the strain from which it originated. These results were further confirmed in I3S. Using a database that contained only the right and left wing samples of all the groups in equal numbers, it was seen that when tested with a right wing, the left wing of the correct strain was just as likely (near 50%) to be ranked early (rank 1 or 2) as the right wing and vice versa for the left wing (Tables 4 & 5).

The test was possible in I3S because the database contained wing images only of the strains being considered – images of other species/strains could not interfere significantly with the selection. Furthermore, unlike identifying images of live individuals, such as whale sharks for which the software was initially developed, it was possible to standardise very accurately how all the wing images were obtained. As with other geometric morphometric techniques, the mosquito wings were photographed using a microscope, all of the images were rapidly aligned the same way using Microsoft Photos – a step which is not always undertaken, but is necessary when considering strains where differences could be minor, but consistent. This permitted the gathering of information in Tables 4 & 5 - the result of imaging dead, therefore still wings under the microscope all aligned the same way, using a systematic method (e.g. coverslip to prevent wing folding). Such levels of standardisation is not possible when taking photographs of live, moving individuals from afar, even though the algorithm takes differences in how the photographs are obtained into consideration when the image is annotated with the required blue dots. Had the right and left wings been different from each other, they would not have selected for each other in the early first and second ranks. Equalising as many variables as possible in the methodology was necessary because the study dealt with closely resembling families and strains (transgenic DSM 1 & 2 derived from *A. gambiae* G3) where the differences would be expected to be small. Taken collectively, all of the results indicated that high levels of bilateral symmetry was present in the wings of DSM 1 & 2 as well as the parent strain *A. gambiae* G3.

Restricting the number of strains and species in the database should not be taken to mean that the correct strain would not be selected from a database containing larger numbers of other species and strains, as DSM1 (from a previous study) was selected 100% of the time at rank 1 from a database containing a total of forty different species and strains and sixty different wing images of mosquito wings (Vyas-Patel N & Mumford JD 2018). It was used here to simplify and equalise the process at every level and to enable a rapid assessment of symmetry between right and left wings, with ease. Also because very small differences (at least by eye) between parents and offspring were being considered.

In addition to wing size and symmetry, wing shape has often been used in studies to detect changes in different populations of a species. It was reported that a change in wing shape was related to the distance of a mosquito population from an urban area and this was probably related to differences in environmental pressures such as chemical control (Wilk-da-Silva R et al, 2018). I3S takes wing shape into account as the wing outline is marked in I3S where the veins meet the edge of the wing. The shape of the wings as well as the cloud of points marking the meeting point of the veins within the wing both contribute to distinguishing the very similar (in appearance) transgenic strains from the parent *A. gambiae* G3 strain in I3S (Vyas-Patel N & Mumford JD, 2018). It is clear from this study that I3S could distinguish between strains of the same species, in this case transgenic DSM 1 & 2 from the parent *A. gambiae* G3 with 70 to 100% accuracy at Rank 1 and could be used in any study where different strains needed to be sorted, provided that a phenotypic difference existed between the strains under consideration. As closely resembling strains were the subjects and I3S sorting can be under 100% (here it was from 70% to 100%) for full assurance, measurements and molecular confirmation for the strain type should also be undertaken.

Obtaining individual measurements of body parts is the norm in studies of larger animals. As soon as an individual is captured and photographed, measurements are taken as a matter of routine procedure, providing valuable information relating to the health of the population. The assessment of size, shape and symmetry of individual insects should also become part of the investigative procedure in all endeavours. Attempts have been made (Müller C, 2018), but it is not part of regular procedure to take the measurements of insects. Partly, this is due to the fact that size is readily measured in larger animals (with a measuring tape and callipers), small insect measurements require more time and effort. In this study, measuring using Microsoft Paint proved to be rapid, easy and provided assurance that the methods being adopted for insect production were proceeding well and had a good chance of success. The transgenic mosquitoes, although phenotypically closely similar to the parents, presented with differences. These differences could be measured using linear measurement and I3S. The transgenic mosquitoes were larger than the population from which they were derived; their wings were bilaterally symmetrical and could be distinguished with a high level of accuracy (70% to 100%) from the parent strain using I3S Classic software.

## Conclusions

- Every newly created organism warrants closer scrutiny as phenotypic differences, however small, may exist and can be measured between closely resembling strains.
- Both sexes of the transgenic mosquitoes DSM1 & 2 were larger than the parent strain *A. gambiae* G3, when wing length and width measurements were assessed. This was a positive indicator of their fitness, especially as they mated with the parent strain when back crossed continually.
- There was no significant wing size difference between DSM1 &2.
- Female wings were always larger (longer and broader) than male wings from all the groups tested.
- Both wing widths and lengths were reliable indicators that could be used to determine the sex of mosquitoes, with male mosquito wings being significantly shorter and narrower than female wings, from all the groups tested. Using wing size, as well as I3S Classic, a clear demarcation of the sexes could be seen from all the groups.
- Transgenic male wings were smaller (shorter and narrower) than the parent *A. gambiae* G3 females as well as the transgenic females.
- Female G3s were larger than the transgenic males DSM1 & 2 as well as male G3s.
- Microsoft Paint and Photos proved to be easy to use tools to obtain accurate measurements and attain image alignment from photos of insect wings. In addition to identifying the correct strain in the early first and second ranks, I3S could also be used to compare right and left wings.
- Of the different methods used to gauge wing symmetry the simplest and equally accurate was to subtract the measurements of the right and left wings and then determine how close to zero those differences were.

## Supporting information

Length data

Width data

## Acknowledgements

With thanks to Imperial College London and Professor John D Mumford for manuscript reading and comment.

