## Supplementary material for "Monitoring transgenic mosquitoes using wing measurements and I3S Classic": Length data

A to L is T1 T2 G3 (right and Left) all F (female), then the same again all M (Male).

**ANOVA: Results**

The results of a ANOVA statistical test performed at 06:51 on 17-OCT-2018

```
 Source of     Sum of       d.f.    Mean         F
 Variation     Squares              Squares

  between      3.878         11    0.3525        25.94
  error        5.545        408    1.3592E-02
  total        9.423        419
```

The probability of this result (‘p’ value), assuming the null hypothesis, is less than .0001

As the p-value is <0.0001, the result is significant at p<0.05.

Group A: Number of items= 35
2.46 2.49 2.66 2.74 2.74 2.75 2.77 2.79 2.81 2.81 2.83 2.84 2.84 2.84 2.85 2.85 2.87 2.88 2.89 2.90 2.90 2.92 2.92 2.92 2.93 2.94 2.96 2.96 2.96 2.96 2.97 3.01 3.03 3.14 3.19

Mean = 2.8663
95% confidence interval for Mean: 2.828 thru 2.905
Standard Deviation = 0.145
High = 3.190 Low = 2.460
Median = 2.880
Average Absolute Deviation from Median = 0.102

Group B: Number of items= 35
2.49 2.51 2.65 2.73 2.73 2.76 2.76 2.78 2.81 2.81 2.83 2.85 2.85 2.85 2.88 2.89 2.89 2.89 2.90 2.90 2.90 2.91 2.93 2.94 2.94 2.94 2.95 2.98 2.99 3.00 3.01 3.03 3.05 3.16 3.22

Mean = 2.8774
95% confidence interval for Mean: 2.839 thru 2.916
Standard Deviation = 0.149
High = 3.220 Low = 2.490
Median = 2.890
Average Absolute Deviation from Median = 0.105

Group C: Number of items= 35
2.61 2.66 2.66 2.72 2.74 2.74 2.75 2.76 2.77 2.79 2.80 2.81 2.82 2.83 2.83 2.83 2.83 2.85 2.85 2.88 2.88 2.88 2.89 2.91 2.91 2.93 2.97 2.98 2.99 2.99 3.00 3.00 3.00 3.11 3.14

Mean = 2.8603
95% confidence interval for Mean: 2.822 thru 2.899
Standard Deviation = 0.122
High = 3.140 Low = 2.610
Median = 2.850
Average Absolute Deviation from Median = 9.600E-02

Group D: Number of items= 35
2.62 2.65 2.67 2.74 2.75 2.76 2.78 2.78 2.79 2.79 2.79 2.79 2.82 2.82 2.82 2.83 2.85 2.86 2.87 2.89 2.91 2.92 2.93 2.95 2.97 2.97 2.98 2.99 2.99 3.01 3.01 3.06 3.10 3.23 3.30

Mean = 2.8854
95% confidence interval for Mean: 2.847 thru 2.924
Standard Deviation = 0.149
High = 3.300 Low = 2.620
Median = 2.860
Average Absolute Deviation from Median = 0.115

Group E: Number of items= 35
2.60 2.66 2.67 2.67 2.68 2.69 2.69 2.71 2.71 2.73 2.73 2.74 2.75 2.76 2.77 2.77 2.77 2.78 2.79 2.79 2.79 2.81 2.81 2.81 2.82 2.82 2.83 2.85 2.87 2.87 2.88 2.89 2.95 2.96 2.97

Mean = 2.7826
95% confidence interval for Mean: 2.744 thru 2.821
Standard Deviation = 8.816E-02
High = 2.970 Low = 2.600
Median = 2.780
Average Absolute Deviation from Median = 6.886E-02

Group F: Number of items= 35
2.61 2.64 2.65 2.66 2.67 2.69 2.69 2.71 2.72 2.72 2.74 2.74 2.74 2.74 2.74 2.75 2.75 2.76 2.78 2.78 2.78 2.79 2.79 2.79 2.80 2.82 2.82 2.83 2.84 2.86 2.87 2.88 2.92 2.98 3.01

Mean = 2.7731
95% confidence interval for Mean: 2.734 thru 2.812
Standard Deviation = 9.006E-02
High = 3.010 Low = 2.610
Median = 2.760
Average Absolute Deviation from Median = 6.800E-02

Group G: Number of items= 35
2.46 2.48 2.49 2.53 2.54 2.56 2.61 2.62 2.63 2.64 2.65 2.66 2.69 2.69 2.69 2.70 2.72 2.72 2.72 2.73 2.74 2.74 2.74 2.75 2.75 2.76 2.76 2.77 2.80 2.80 2.81 2.82 2.82 2.83 2.85

Mean = 2.6934
95% confidence interval for Mean: 2.655 thru 2.732
Standard Deviation = 0.105
High = 2.850 Low = 2.460
Median = 2.720
Average Absolute Deviation from Median = 8.086E-02

Group H: Number of items= 35
2.34 2.44 2.46 2.49 2.53 2.55 2.61 2.61 2.63 2.65 2.65 2.66 2.66 2.69 2.71 2.72 2.72 2.73 2.73 2.73 2.73 2.74 2.75 2.76 2.76 2.76 2.77 2.78 2.79 2.79 2.80 2.80 2.82 2.86 2.87

Mean = 2.6883
95% confidence interval for Mean: 2.650 thru 2.727
Standard Deviation = 0.123
High = 2.870 Low = 2.340
Median = 2.730
Average Absolute Deviation from Median = 8.914E-02

Group I: Number of items= 35
2.45 2.48 2.50 2.53 2.54 2.55 2.56 2.58 2.58 2.61 2.63 2.64 2.64 2.64 2.65 2.66 2.67 2.68 2.68 2.68 2.69 2.69 2.69 2.69 2.70 2.71 2.73 2.74 2.75 2.76 2.76 2.77 2.77 2.79 2.80

Mean = 2.6569
95% confidence interval for Mean: 2.618 thru 2.696
Standard Deviation = 9.103E-02
High = 2.800 Low = 2.450
Median = 2.680
Average Absolute Deviation from Median = 7.114E-02

Group J: Number of items= 35
2.45 2.47 2.52 2.53 2.55 2.56 2.56 2.58 2.59 2.62 2.64 2.65 2.65 2.66 2.66 2.66 2.66 2.67 2.67 2.67 2.68 2.69 2.69 2.69 2.69 2.71 2.71 2.71 2.72 2.73 2.74 2.76 2.79 2.79 2.79

Mean = 2.6546
95% confidence interval for Mean: 2.616 thru 2.693
Standard Deviation = 8.552E-02
High = 2.790 Low = 2.450
Median = 2.670
Average Absolute Deviation from Median = 6.343E-02

Group K: Number of items= 35
2.42 2.43 2.45 2.49 2.49 2.53 2.53 2.56 2.57 2.58 2.59 2.63 2.63 2.63 2.63 2.63 2.65 2.65 2.65 2.67 2.67 2.69 2.69 2.71 2.72 2.72 2.73 2.74 2.74 2.75 2.77 2.77 2.78 2.82 2.82

Mean = 2.6437
95% confidence interval for Mean: 2.605 thru 2.682
Standard Deviation = 0.108
High = 2.820 Low = 2.420
Median = 2.650
Average Absolute Deviation from Median = 8.571E-02

Group L: Number of items= 35
2.41 2.43 2.43 2.46 2.46 2.51 2.52 2.53 2.57 2.59 2.59 2.61 2.61 2.61 2.62 2.63 2.63 2.63 2.64 2.66 2.68 2.69 2.69 2.69 2.72 2.72 2.72 2.74 2.75 2.75 2.76 2.79 2.80 2.82 2.82

Mean = 2.6366
95% confidence interval for Mean: 2.598 thru 2.675
Standard Deviation = 0.115
High = 2.820 Low = 2.410
Median = 2.630
Average Absolute Deviation from Median = 9.229E-02

**Data Reference: 0EEE**

**Plot the Group Means with 95% Confidence Intervals**

Top of Form

**Format:**
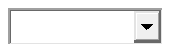

**Y Scale Options:**

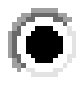
Linear: base near ymin

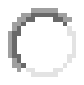
Linear: base y=0

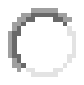
Log

**Point Symbol Options:**

The symbol should be
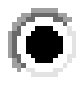
solid (filled) or
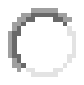
outline only

Type of symbol:

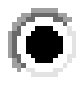
Circles

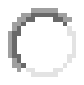
Squares

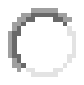
Diamonds

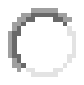
Triangles Up

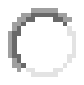
Triangles Down

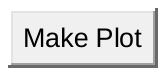

Bottom of Form

**Box Plot the Data**

Top of Form

**Format:**
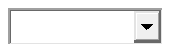

**Y Scale Options:**

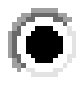
Linear

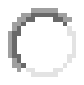
Log

**Options:**

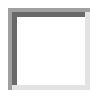
data swarm

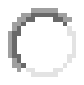
mean with 1
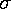
 error bars

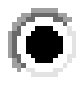
boxplot

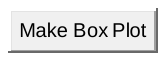

Bottom of Form
