## Supplementary material for "Monitoring transgenic mosquitoes using wing measurements and I3S Classic": Width data

ANOVA Widths-

**ANOVA: Results**

The results of a ANOVA statistical test performed at 09:27 on 20-OCT-2018

```
 Source of     Sum of       d.f.    Mean         F
 Variation     Squares              Squares

  between      1.215         11    0.1104        110.0
  error       0.4096        408    1.0039E-03
  total        1.624        419
```

The probability of this result, assuming the null hypothesis, is less than .0001

Group A: Number of items= 35
0.680 0.680 0.690 0.690 0.690 0.690 0.690 0.690 0.700 0.710 0.710 0.710 0.710 0.720 0.720 0.720 0.730 0.730 0.740 0.740 0.740 0.740 0.740 0.740 0.750 0.750 0.750 0.750 0.750 0.760 0.760 0.760 0.770 0.780 0.790

Mean = 0.72771
95% confidence interval for Mean: 0.7172 thru 0.7382
Standard Deviation = 2.981E-02
High = 0.7900 Low = 0.6800
Median = 0.7300
Average Absolute Deviation from Median = 2.514E-02

Group B: Number of items= 35
0.630 0.670 0.680 0.680 0.680 0.690 0.690 0.700 0.710 0.710 0.710 0.720 0.720 0.720 0.730 0.730 0.730 0.730 0.740 0.740 0.740 0.750 0.750 0.750 0.750 0.750 0.750 0.750 0.760 0.760 0.760 0.760 0.770 0.780 0.790

Mean = 0.72800
95% confidence interval for Mean: 0.7175 thru 0.7385
Standard Deviation = 3.471E-02
High = 0.7900 Low = 0.6300
Median = 0.7300
Average Absolute Deviation from Median = 2.714E-02

Group C: Number of items= 35
0.670 0.680 0.680 0.680 0.690 0.690 0.690 0.700 0.710 0.710 0.710 0.710 0.710 0.720 0.720 0.720 0.720 0.720 0.730 0.730 0.740 0.740 0.740 0.740 0.750 0.750 0.750 0.750 0.750 0.750 0.760 0.770 0.780 0.790 0.790

Mean = 0.72686
95% confidence interval for Mean: 0.7163 thru 0.7374
Standard Deviation = 3.160E-02
High = 0.7900 Low = 0.6700
Median = 0.7200
Average Absolute Deviation from Median = 2.571E-02

Group D: Number of items= 35
0.650 0.670 0.690 0.690 0.690 0.690 0.690 0.700 0.710 0.710 0.710 0.710 0.710 0.720 0.720 0.720 0.730 0.730 0.730 0.740 0.740 0.740 0.740 0.740 0.750 0.760 0.760 0.760 0.760 0.770 0.770 0.780 0.790 0.820 0.880

Mean = 0.73343
95% confidence interval for Mean: 0.7229 thru 0.7440
Standard Deviation = 4.379E-02
High = 0.8800 Low = 0.6500
Median = 0.7300
Average Absolute Deviation from Median = 3.200E-02

Group E: Number of items= 35
0.650 0.650 0.660 0.670 0.680 0.690 0.690 0.690 0.690 0.690 0.690 0.700 0.710 0.710 0.710 0.710 0.710 0.720 0.720 0.720 0.720 0.730 0.730 0.730 0.730 0.730 0.730 0.730 0.730 0.740 0.740 0.750 0.750 0.760 0.770

Mean = 0.71229
95% confidence interval for Mean: 0.7018 thru 0.7228
Standard Deviation = 2.931E-02
High = 0.7700 Low = 0.6500
Median = 0.7200
Average Absolute Deviation from Median = 2.314E-02

Group F: Number of items= 35
0.630 0.660 0.670 0.670 0.680 0.680 0.680 0.690 0.690 0.690 0.690 0.690 0.690 0.690 0.700 0.700 0.700 0.700 0.700 0.710 0.710 0.710 0.720 0.730 0.730 0.740 0.740 0.740 0.740 0.750 0.750 0.760 0.760 0.780 0.780

Mean = 0.71000
95% confidence interval for Mean: 0.6995 thru 0.7205
Standard Deviation = 3.456E-02
High = 0.7800 Low = 0.6300
Median = 0.7000
Average Absolute Deviation from Median = 2.714E-02

Group G: Number of items= 35
0.550 0.570 0.570 0.590 0.590 0.590 0.610 0.610 0.620 0.620 0.620 0.620 0.620 0.630 0.630 0.630 0.630 0.630 0.630 0.640 0.640 0.640 0.640 0.640 0.640 0.640 0.640 0.650 0.650 0.650 0.650 0.650 0.650 0.680 0.680

Mean = 0.62686
95% confidence interval for Mean: 0.6163 thru 0.6374
Standard Deviation = 2.847E-02
High = 0.6800 Low = 0.5500
Median = 0.6300
Average Absolute Deviation from Median = 2.029E-02

Group H: Number of items= 35
0.530 0.550 0.560 0.570 0.590 0.590 0.610 0.610 0.610 0.620 0.620 0.620 0.620 0.620 0.630 0.630 0.630 0.630 0.630 0.630 0.630 0.640 0.640 0.640 0.640 0.640 0.640 0.640 0.640 0.640 0.640 0.650 0.650 0.650 0.660

Mean = 0.62114
95% confidence interval for Mean: 0.6106 thru 0.6317
Standard Deviation = 2.978E-02
High = 0.6600 Low = 0.5300
Median = 0.6300
Average Absolute Deviation from Median = 1.971E-02

Group I: Number of items= 35
0.560 0.560 0.580 0.580 0.580 0.590 0.590 0.590 0.590 0.590 0.600 0.600 0.600 0.610 0.610 0.610 0.620 0.620 0.620 0.620 0.630 0.630 0.630 0.630 0.630 0.630 0.640 0.640 0.640 0.640 0.650 0.650 0.650 0.650 0.650

Mean = 0.61457
95% confidence interval for Mean: 0.6040 thru 0.6251
Standard Deviation = 2.638E-02
High = 0.6500 Low = 0.5600
Median = 0.6200
Average Absolute Deviation from Median = 2.200E-02

Group J: Number of items= 35
0.560 0.560 0.570 0.580 0.580 0.590 0.590 0.590 0.590 0.590 0.590 0.590 0.600 0.600 0.600 0.610 0.610 0.610 0.610 0.610 0.620 0.620 0.620 0.620 0.620 0.630 0.630 0.630 0.640 0.640 0.640 0.640 0.640 0.650 0.650

Mean = 0.60914
95% confidence interval for Mean: 0.5986 thru 0.6197
Standard Deviation = 2.478E-02
High = 0.6500 Low = 0.5600
Median = 0.6100
Average Absolute Deviation from Median = 2.029E-02

Group K: Number of items= 35
0.550 0.550 0.570 0.570 0.580 0.580 0.590 0.590 0.590 0.590 0.590 0.590 0.590 0.600 0.600 0.610 0.610 0.620 0.620 0.620 0.620 0.620 0.620 0.630 0.640 0.640 0.640 0.640 0.650 0.650 0.650 0.650 0.650 0.670 0.670

Mean = 0.61286
95% confidence interval for Mean: 0.6023 thru 0.6234
Standard Deviation = 3.195E-02
High = 0.6700 Low = 0.5500
Median = 0.6200
Average Absolute Deviation from Median = 2.657E-02

Group L: Number of items= 35
0.550 0.560 0.570 0.570 0.580 0.580 0.590 0.590 0.590 0.590 0.600 0.600 0.600 0.600 0.610 0.610 0.610 0.620 0.620 0.620 0.620 0.620 0.630 0.630 0.640 0.640 0.640 0.640 0.650 0.650 0.650 0.650 0.650 0.670 0.670

Mean = 0.61457
95% confidence interval for Mean: 0.6040 thru 0.6251
Standard Deviation = 3.090E-02
High = 0.6700 Low = 0.5500
Median = 0.6200
Average Absolute Deviation from Median = 2.543E-02

**Data Reference: 717D**

**Plot the Group Means with 95% Confidence Intervals**

Top of Form

**Format:**
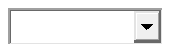

**Y Scale Options:**

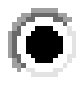
Linear: base near ymin

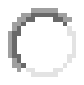
Linear: base y=0

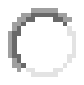
Log

**Point Symbol Options:**

The symbol should be
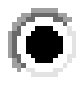
solid (filled) or
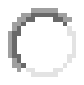
outline only

Type of symbol:

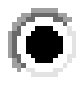
Circles

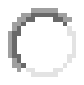
Squares

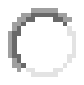
Diamonds

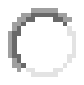
Triangles Up

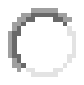
Triangles Down

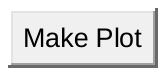

Bottom of Form

**Box Plot the Data**

Top of Form

**Format:**
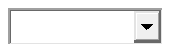

**Y Scale Options:**

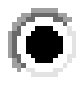
Linear

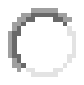
Log

**Options:**

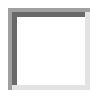
data swarm

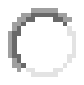
mean with 1
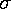
 error bars

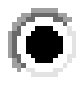
boxplot

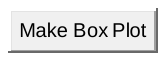

Bottom of Form
